# Reactive Oxygen Species Counteract Wound Contraction and Promote Wound Healing

**DOI:** 10.1101/2025.03.27.645726

**Authors:** Chang Ding, Linlin Li, Yueyang Wang, Hong-Anh A. Nguyen, Deva D. Chan, David M. Umulis, Adrian T. Buganza, Qing Deng

**Affiliations:** Department of Biological Sciences, Purdue University, West Lafayette, IN 47907, USA; Weldon School of Biomedical Engineering, Purdue University, West Lafayette, IN47907, USA

**Keywords:** reactive oxygen species (ROS), amputation wound, actomyosin, linear elasticity simulation, zebrafish

## Abstract

Reactive oxygen species (ROS) are second messengers that drive wound closure. However, the mechanism by which ROS regulate wound contraction to facilitate wound healing remains unclear. Here, we report that ROS counteract wound contraction by inhibiting the phosphorylation of myosin regulatory light chain. Acute ROS inhibition, through pharmacological perturbations, disturbs wound relaxation, delays wound closure, and impairs regrowth following amputation. Moreover, actomyosin inhibition relaxes tailfin contraction without impairing wound closure or regrowth. Over-contraction, on the other hand, impedes wound closure. Meanwhile, chronic depletion of epithelial ROS during embryonic development, achieved through morpholino-mediated knockdown of the *duox* gene, alters tissue stiffness, as measured using atomic force microscopy-based nanoindentation. Despite a reduced contraction force, the wound also appears to be over-contracted, with delayed healing and regrowth. An *in silico* linear elasticity simulation to calculate the second principal stress based on node-wise prescribed displacement recapitulated the contraction dynamics during acute and chronic ROS inhibition. Together, our results provide a novel understanding of how reactive oxygen species (ROS) facilitate wound closure, a process instrumental in restoring tissue integrity and maintaining homeostasis.

## INTRODUCTION

Wound closure is essential for reestablishing epithelia’s barrier function after injury (Redd *et al*., 2004). Epithelial wounds in aquatic environments, such as those in the zebrafish tailfin, and injuries in the mucosal lining of the upper respiratory and digestive tracts in mammals are exposed to excessively high microbial burdens in environmental fluids (Naomi *et al*., 2021). An efficient wound repair mechanism is essential for the survival of aquatic species and the homeostasis at mucosal surfaces.

Upon wounding, multiple signals, including electric potential differences, reduced junction signals, osmolarity differences, and chemical signals, coordinate to close the wound (Gault, Enyedi and Niethammer, 2014; Enyedi and Niethammer, 2015; JIA *et al*., 2021). Most wound signals converge on the second messenger, calcium. ROS are other second messengers that facilitate wound healing in various species, including drosophila and zebrafish (Razzell *et al*., 2013; de Oliveira *et al*., 2014). In both systems, calcium triggers H_2_O_2_ accumulation at the wound edge within 10 mins post-amputation (mpa) through the activation of NADPH Oxidase (NOX) (Niethammer *et al*., 2009; Razzell *et al*., 2013; de Oliveira *et al*., 2014).

ROS, including superoxide, hydroxyl radical, hydrogen peroxide (H_2_O_2_), and singlet oxygen, are redox-sensitive species that orchestrate complex physiological signaling processes (Sies *et al*., 2022). Among all ROS species, H_2_O_2_ is the most relevant in conveying long-range signals due to its longer half-life, higher permeability, and significantly reduced cellular toxicity compared to other ROS species (Finkel, 1998; Steiling *et al*., 1999; Kanta, 2011). H_2_O_2_ facilitates wound healing by mediating various wound-signaling pathways (Basuroy *et al*., 2010; Vatsyayan *et al*., 2011; Soares *et al*., 2016; David *et al*., 2017). Firstly, H_2_O_2_ regulates the activation of the transcription factor nuclear factor erythroid-derived 2-like 2 (Nrf2), promoting re-epithelialization. (Soares *et al*., 2016; David *et al*., 2017). Secondly, H_2_O_2_ leads to lipid peroxidation, which facilitates cell proliferation (Vatsyayan *et al*., 2011). Upon wounding, a low level of 4-hydroxynonenal (4-HNE) is synthesized through the oxidation of H_2_O_2_, which phosphorylates the epidermal growth factor receptor (EGFR) and activates downstream proliferation-related kinases, such as Akt and ERK 1/2, to promote wound healing. Thirdly, H_2_O_2_ may directly modulate kinase activities, such as phosphorylating FAK kinase, to facilitate cell adhesion and migration during wound closure (Basuroy *et al*., 2010). Fourthly, H_2_O_2_ can directly or indirectly regulate actin cytoskeleton dynamics, either through the direct oxidation of actin or the regulation of gene expression of actin-binding proteins (ABPs), including Profilin, ARP-3, MRLC, and 14-3-3ζ, thereby inducing wound closure (Clarkson *et al*., 2002). It is worth noting that the H_2_O_2_ level is tightly regulated during wound-healing. Excessive H_2_O_2_ level elevates inflammatory transcription factor activation, such as activator protein 1 (AP-1) and nuclear factor kappa B (NF-ĸB), leading to extracellular matrix protein degradation and delayed wound healing (Schäfer and Werner, 2008; Dunnill *et al*., 2017). A too low level of H_2_O_2_ also impedes wound healing by impairing fibroblast functions (Fujiwara *et al*., 2016). However, mechanisms underlying how H_2_O_2_ promotes mucosal wound healing in multicellular tissue remain to be further elucidated, mainly due to the complexity of the signaling networks.

The zebrafish larvae tailfin model has been utilized to characterize the role of H_2_O_2_ in mucosal wound healing. It mimics a similar microbial selection pressure in the surrounding aquatic environments (Naomi *et al*., 2021). In addition, the high regenerative ability and transparency of the tissue made it an ideal model for studying wound closure and tissue regeneration (Gemberling *et al*., 2013). In this model, H_2_O_2_ is primarily generated from the enzyme Dual oxidase (Duox), an isoform of NADPH oxidase (Mitchison *et al*., 2009; Razzell *et al*., 2013). Mulero *et al*. demonstrated that calcium activates Duox and produces H_2_O_2_ to trigger NF-kB signaling through the p2yR/PLC/calcium signaling pathway for wound closure (de Oliveira *et al*., 2014). Similarly, injury-generated H_2_O_2_ at the tailfin induces Fynb activation, a family member of Src family kinase, for wound regeneration (Yoo *et al*., 2012). Although previous studies have suggested that H_2_O_2_ plays a pivotal role in wound closure, the understanding of how H_2_O_2_ regulates cytoskeleton compartments to seal the wound remains limited.

Here, we report a novel perspective on the function of ROS in facilitating wound healing, whereby ROS relax the wound mechanically to promote wound closure. The zebrafish fin fold amputated wound contracts for 20-30 mins post-wounding, and then it gradually relaxes to its initial width (Gault, Enyedi and Niethammer, 2014). The formation of the actomyosin ring provides the main driving force for wound contraction and has long been regarded as beneficial for wound closure (Abreu-Blanco et al., 2012; Clark et al., 2009; Russo et al., 2005). Surprisingly, we found that the contraction is disposable for the closure of amputation wounds. ROS relaxes wounds to prevent over-contraction, which impedes wound closure. Together, our work reveals that ROS promote relaxation and closure of amputation wound, demonstrating a context-dependent role of actomyosin in mediating mucosal wound closure, which is instrumental in our understanding of wound biology.

## RESULTS

### ROS promote wound relaxation

To address the function of ROS in regulating wound contraction, we incorporate a NOX inhibitor diphenylene iodonium (DPI), a ROS scavenger N-acetyl-cysteine (NAC), and *duox* morpholino (MO) to eliminate ROS (Fig. 1). Pretreatment of DPI and NAC significantly reduced H_2_O_2_ level at the wound edge at 20 min post-amputation (mpa), as reflected by the increase in pentafluorobenzenesulfonyl fluoresce, which has a higher selectivity of H_2_O_2_ over other ROS species (Fig. 1A, B). Consistent with previous observations, the DMSO control contract for 15 mpa and gradually relaxed to its initial tailfin width 1h after wounding (Gault et al., 2014). However, the tailfin of both DPI-treated and NAC-treated larvae continued to contract without returning to the relaxed tailfin width (Fig. 1C-E and Movie 1). Similarly, *duox* morphants were defective in pre-mRNA splicing and displayed significantly reduced H_2_O_2_ production at the wound edge (Fig. 1F-H). In addition, the *duox* morphants displayed more robust and prolonged wound contraction compared to the *p53* morphant control (Fig. 1I, J, and Movie 2). Together, ROS promote wound relaxation after amputation.

**Figure 1.**
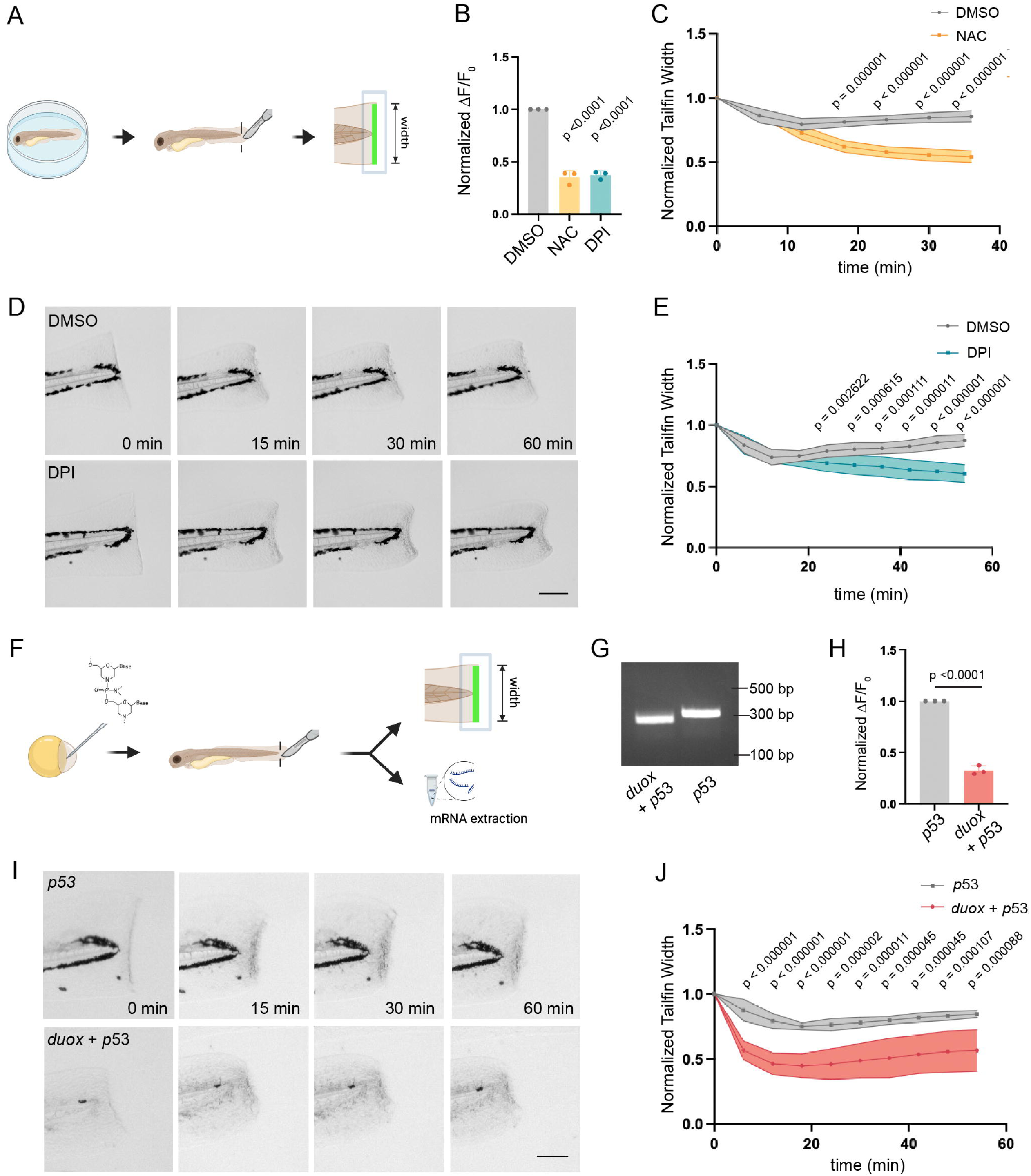
ROS drive wound relaxation. (A) Workflow of zebrafish embryo fin fold amputation. Embryos at 3 dpf were amputated just posterior to the notochord. ROS intensity was quantified at the wound edge. The width of the wound is plotted over time to show wound contraction and relaxation. (B) Relative ROS production at wound edge of NAC-treated, DPI-treated, and DMSO ctrl groups. Fish were treated for 1h before wounding and maintained in inhibitor for 1h post wounding. ROS production at 20 post-amputation (mpa) was normalized to 0 mpa for each group. Data are presented as mean ± s.e.m. from three independent experiments, with n = 20 larvae per group in each experiment. (C) Wound widths at indicated time points were normalized to 0 mpa for the NAC-treated or DMSO ctrl group. n=9. Data are presented as mean ± s.e.m., pooled from three independent experiments. (D) Representative images of wound dynamics in the DPI-treated and DMSO ctrl group at 0, 15, 30, and 60 mpa. Scale bars: 50 µm. (E) Wound widths at the indicated time points were normalized to 0 mpa for the DPI-treated (n = 9) and DMSO ctrl (n = 10) groups. Data are mean ± s.e.m. pooled from 3 independent experiments. (F) Workflow of zebrafish embryo fin fold amputation with *duox* knockdown. One-cell stage embryos were injected with either *duox*+*p53* or *p53* morpholino. Embryos at 3 dpf were wounded as described in A. In addition, mRNA was collected to validate the *duox* knockdown. (G) RT-PCR was used to confirm the MO-induced splicing defect. (H) Relative ROS production at wound edge of *p53* –injected ctrl and *duox + p53*-injected groups. Data are presented as mean ± s.e.m. from three independent experiments, with n = 15 larvae per group in each experiment. (I) Representative images of wound dynamics in the *duox + p53-*injected group and the *p53-*injected ctrl group at 0, 15, 30, and 60 mpa. Scale bars: 50 µm. (J) Wound widths at indicated time points were normalized to 0 mpa for *duox + p53-*injected (n = 9) and *p53-injected* ctrl (n = 9) groups. Data are presented as mean ± s.e.m., pooled from four independent experiments. (B) One-Way ANOVA. (C, E, H, J) Unpaired *t*-test. Schematics in (A) and (F) were created using BioRender.com.

### ROS are required for wound closure and regrowth

To determine whether ROS regulate wound closure, we utilized a transgenic line *Tg(krt4: NLS-GFP)^pu50^* to quantify the wound opening by measuring the distances of nuclei of the enveloping epithelial layer to the rendered wound center at the wound edge (Fig. 2A). We observed reduced wound opening is at 30 mpa compared with 0 mpa (Fig. 2B, C, E), consistent with the previous work that an amputated wound takes approximately 30 min to seal. (Franco *et al*., 2019). In contrast, DPI treatment or *duox* knockdown expanded the wound opening at 30 mpa (Fig. 2B-F). We measured regrowth after amputation to further determine the physiological consequence (Fig. 2G). DPI was washed out at 1 hpa, to prevent prolonged ROS inhibition. Again, DPI treatment or *duox* knockdown significantly impaired regrowth compared to their control groups at 2 days post-amputation (dpa), consistent with the wound closure defect (Fig. 2H, I). Furthermore, *duox* knockdown increased mortality at 2 dpa, whereas no mortality was observed in the controls (Fig. 2J). Together, ROS are required for rapid wound healing and subsequent regrowth and survival after amputation.

**Figure 2.**
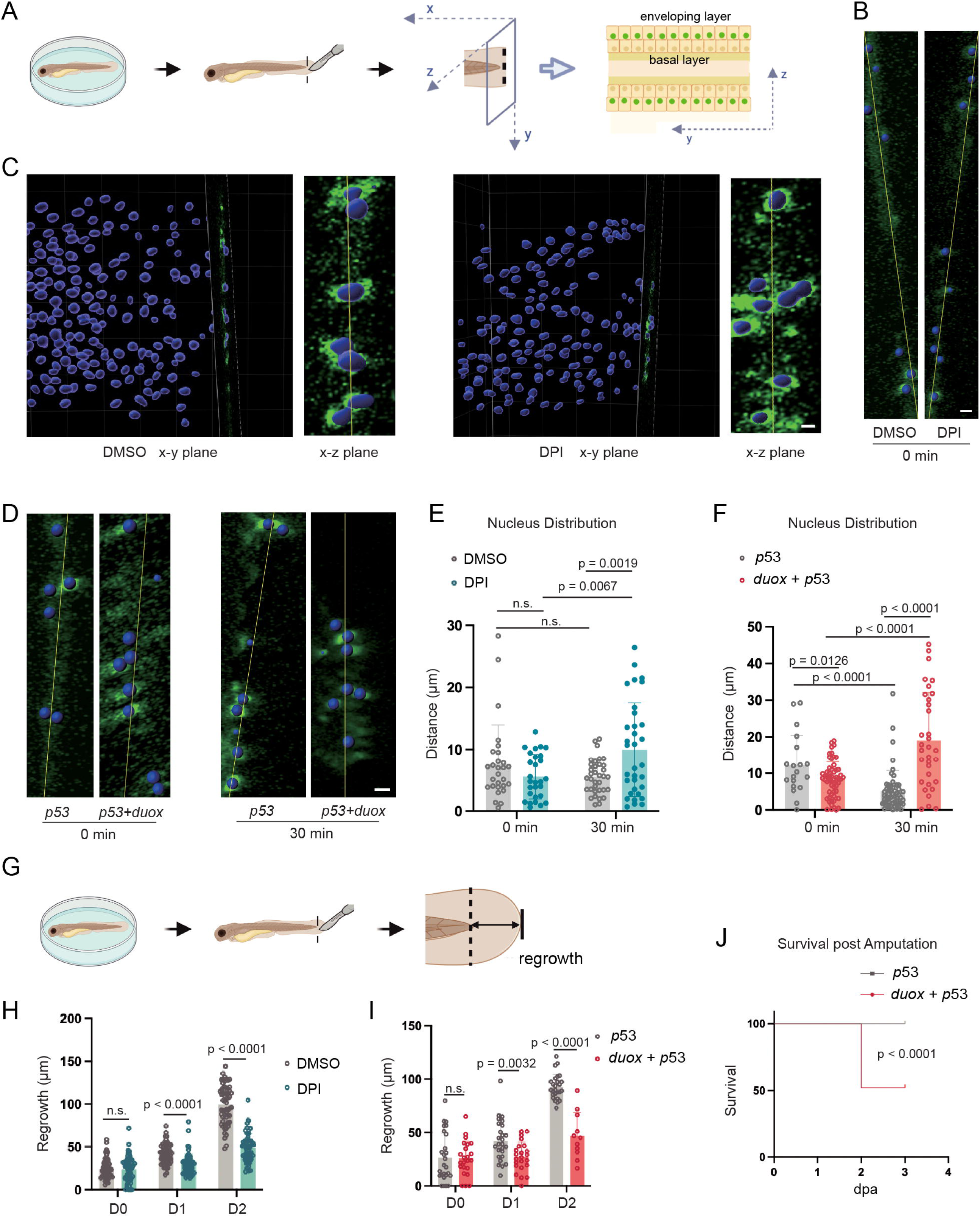
ROS promote wound closure and fin regrowth. (A) Workflow of the zebrafish embryo fin-fold wound closure assay. The nuclei of the enveloping layer are labeled with GFP, and their distance to the wound center is measured at 30 mpa. (B) Representative images of the x-z plane of the wound edge of DPI-treated and DMSO ctrl groups at 0 mpw. Scale bar: 10 µm. (C) Representative images of indicated planes of the wound edge of DPI-treated and DMSO groups at 30 mpa. Scale bar: 5 µm. (D) Representative images of the x-z plane of the wound edge of duox + p53-injected and p53-injected ctrl at 0 and 30 mpa. Blue indicates nuclei masks in (B-D). (E, F) Distances of nuclei at the wound edge to the wound center of inhibitor-treated or morpholino-treated groups at 0 and 30 mpa. Data are presented as mean ± s.e.m., pooled from two independent experiments (n > 20). Scale bar: 10 µm. (G) Workflow of zebrafish embryo fin fold regrowth assay. The distances between the notochord and the fin tip were measured at 0-, 1-, and 2-days pa. (H) Regrowth of DPI-treated and DMSO ctrl groups at 0-, 1-, and 2-days pa. Data are presented as mean ± s.e.m., pooled from two independent experiments (n = 60). (I) Regrowth of *duox* morpholino-treated (n=23) and *p53* ctrl (n=26) groups at 0-, 1-, and 2-days pa. Data are presented as mean ± s.e.m., pooled from two independent experiments (n > 20). (J) Kaplan-Meier survival chart of morpholino-treated groups post-amputation. Data are pooled from 3 independent experiments (n > 20). (E, F, H, I) Two-Way ANOVA. (J) Log-rank analysis. Schematics in (A) and (G) were created using BioRender.com.

### ROS suppress myosin regulatory light chain II phosphorylation

Both F-actin and myosin accumulate at the wound margin as early as 5 mpa, and the actomyosin ring diminishes when the wound fully returns to the relaxed state in 2 dpf larva (Mateus et al., 2012). Since actomyosin is the primary driving force for wound contraction and closure, we next determined how ROS regulate its activity by staining phosphorylated myosin light chain II (p-MRLC) and F-actin (Abreu-Blanco et al., 2012; Clark et al., 2009; Russo et al., 2005) (Fig. 3A-B). Indeed, the p-MRLC and F-actin levels are significantly elevated at 15 mpa compared to 2 mpa (Fig. 3C-E). More importantly, DPI treatment significantly elevated the p-MRLC and F-actin levels at the wound edge compared to the DMSO control at 15 mpa, in line with enhanced wound contraction. Therefore, ROS presumably reduce actomyosin contraction activity, allowing wounds to relax.

**Figure 3.**
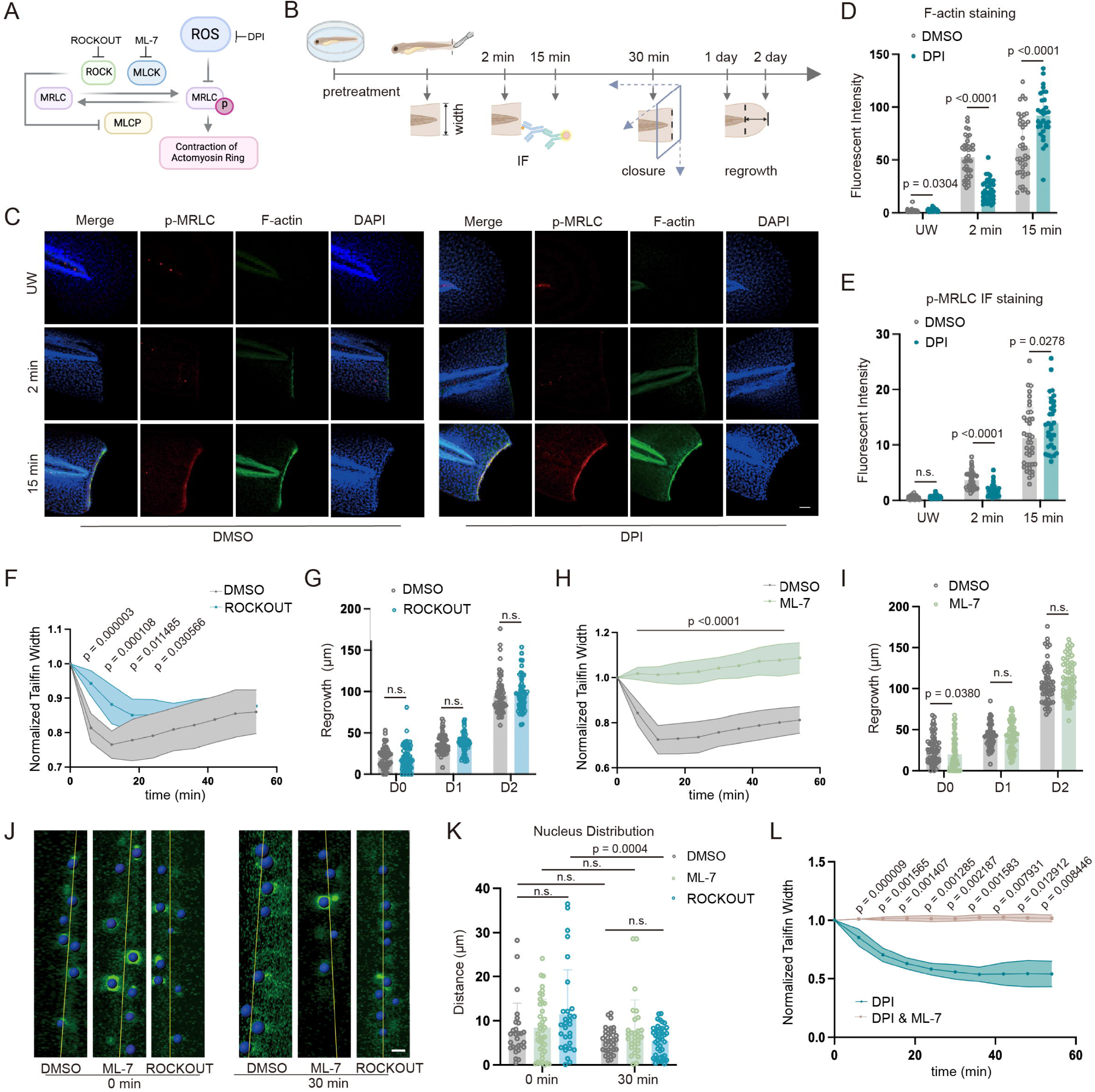
ROS inhibit MRLC phosphorylation and F-actin. (A) Proposed mechanism. ROS inhibit the phosphorylation of MRLC and reduce actin contraction. ROCKOUT inhibits ROCK, and ML-7 inhibits MLCK. (B) Schematic of wound dynamics, closure, and regrowth assay for ROCKOUT, ML-7-treated, and DMSO ctrl groups and immunofluorescence staining assays for DPI-treated and DMSO ctrl groups (C) Representative images of immunofluorescent staining of p-MRLC for DPI-treated and DMSO ctrl groups unwounded (UW) and at 2 min and 15 min pa. Embryos were also stained with phalloidin and DAPI to reveal F-actin and the nucleus. Scale bar: 50 µm. (D, E) p-MRLC and F-actin intensity of DPI-treated and DMSO ctrl groups at the indicated time points. are presented as mean ± s.e.m., pooled from two independent experiments (n > 30). (F) Changes in normalized tailfin width over time in the ROCKOUT-treated group and DMSO ctrl group. are presented as mean ± s.e.m., pooled from three independent experiments (n = 9). (G) Regrowth of ROCKOUT-treated and DMSO ctrl at 0-, 1-, and 2-days pa. Data are presented as mean ± s.e.m., pooled from three independent experiments (n = 60). (H) Changes in normalized tailfin width over time of the ML-7-treated and DMSO ctrl groups. Data are presented as mean ± s.e.m., pooled from three independent experiments (n = 9). (I) Regrowth of ML-7 and ROCKOUT-treated groups and DMSO ctrl group at 0-, 1-, and 2-days pa. Data are presented as mean ± s.e.m., pooled from three independent experiments (n = 60). (J) Representative images of ROCKOUT, ML-7-treated, and DMSO ctrl group’s x-z plane amputated wounds at 0 and 30 mpa. Blue, segmented nuclei. (K) Changes in normalized tailfin width over time of ML-7-treated (n = 9) and ML-7, DPI double-treated groups (n = 10). Data are presented as mean ± s.e.m., pooled from three independent experiments. Scale bar: 10 µm. (L) Relative distances of nuclei at the wound edge to the central plane of the wound in ML-7-treated, ROCKOUT-treated, and DMSO ctrl groups at 0 and 30 mpa. Data are presented as mean ± s.e.m., pooled from two independent experiments (n > 20). (F, H, J) Unpaired *t*-test. (D, E, G, I, L) Two-Way ANOVA. Schematics in (A) and (B) were created using BioRender.com.

### Contraction is not required for wound closure

The Rho-Kinase (ROCK) and Myosin Light Chain Kinase (MLCK) are the two kinases responsible for MRLCII phosphorylation. We then employed ROCK inhibitor ROCKOUT and MLCK inhibitor ML-7 to inhibit p-MLRC, confirming that wound contraction is indeed a result of actomyosin activation (Fig. 3A, F, H). ROCKOUT treatment reduced initial contraction, whereas ML-7 completely blocked wound contraction (Fig. 3A, F, H, and Movie 3). Surprisingly, neither ROCKOUT nor ML-7 treatment impaired wound closure or regrowth (Fig. 3G, I-K), suggesting that wound closure is separable from wound contraction. In addition, ML-7 prevented DPI-induced over-contraction (Fig. 3L), indicating that ROS deficiency-induced over-contraction is dependent on p-MLRC activity.

### Long-term depletion of ROS impairs myosin distribution and reduces tissue stiffness

We next determined whether chronic ROS depletion, using the *duox* morpholino, caused a similar increase in p-MLRC. Despite the prolonged contraction, *duox* morphants had reduced p-MRLC and F-actin levels at the wound edge compared to the *p53* ctrl group at 2 and 15 mpa (Fig. 4A-E). Given that *duox* morphants display a different morphology than the p53 control, with shorter and thinner tailfins, we speculated that the epithelium morphology and its cytoskeleton were perturbed before wounding. The MRLC distribution and cell morphology, revealed by the *Tg(krt4: mrlc2-mScarlet)^pu51^* labeling and plasma membrane stain, in the suprabasal layer of *duox* morphants is altered (Fig. 4F-G). Unlike the uniformly distributed MRLCII at the perijunctional actomyosin ring in the *p53* control group (Tamada *et al*., 2007), the MRLCII in *duox* morphants was scattered. The microridge, revealed by the membrane stain, was abolished. Furthermore, *duox* morphants exhibit altered cell morphology, with a significantly decreased aspect ratio and size compared to the p53 control group (Fig. 4G).

**Figure 4.**
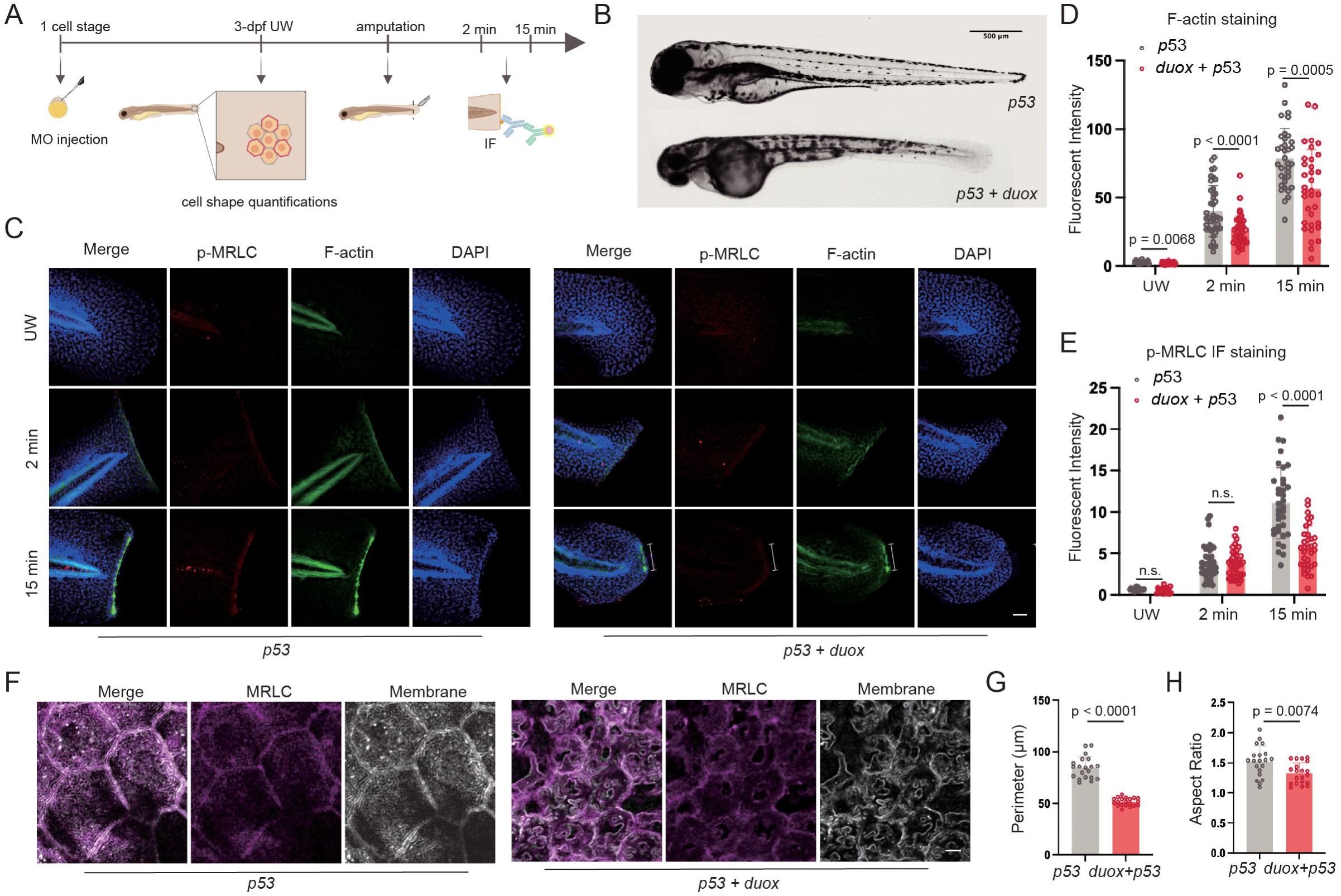
Long-term depletion of ROS impairs myosin activation and epithelial cell morphology. (A) Workflow of MRLC distribution and cell morphology quantification before amputation and immunofluorescence staining of p-MRLC post-amputation in MO-injected groups. (B) Representative morphology of the *duox* morphant and *p53* ctrl groups. Scale bars: 500 µm. (C) Representative images of immunofluorescence of p-MRLC for MO-injected groups unwounded (UW) or at 2 min and 15 min pa. Embryos were also stained with phalloidin and DAPI to reveal F-actin and the nucleus. Scale bar: 50 µm. (D, E) F-actin and p-MRLC intensity of morpholino-injected groups at indicated time points. Data are presented as mean ± s.e.m., pooled from two independent experiments (n>30). (F) Representative images of MRLC and plasma membrane in suprabasal epithelia for MO-injected groups. Scale bar: 5 µm. (G, H) Quantification of the suprabasal epithelial cell membrane morphology for MO-injected groups. Data are presented as mean ± s.e.m., pooled from three independent experiments (n = 20). (D, E) Two-Way ANOVA. (G, H) Unpaired *t*-test. Schematics in (A) were created using BioRender.com.

### Distinct mechanisms contribute to wound over-contraction upon acute and chronic ROS depletion

These developmental defects in *duox* morphants may contribute to the dynamics of wound contraction. We measured the elastic modulus of the *duox* and *p53* morphants using atomic force microscopy (AFM) based nanoindentation. As expected, *duox* morphants had a significantly lower tissue rigidity (Fig. 5A-B). Therefore, we hypothesized that reduced tissue rigidity is the primary reason driving over-contraction in *duox* morphants despite reduced contraction force (Fig. 4E). To examine this hypothesis, we performed an *in silico* linear elasticity simulation to calculate the second principal stress based on the node-wise prescribed displacement to reflect the compression stress at the amputated wound margin for the MO-injected groups (Fig. 1I, J, Fig. 5C-F). The *in silico* wound margin tip displacement was fitted to the experimental data through Fourier transformation (Fig. 1D, Fig. S1). The maximum compressive principal stress of the *p53* control continues to increase as the wound contracts. It peaks at approximately 1000s post-amputation and gradually diminishes as the wound relaxes back to its initial tailfin width (Fig. 5E and Movie 4). The midrange of elastic modulus for each group was used to simulate the principal stress at the wound margin, which was 45175 Pa or 10942 Pa. (Fig. 5B). The maximum stress at the wound edge in the *p53* and *duox* morphants was estimated to be 11386.99 N/m² and 6207.64 N/m², respectively. The significant increase in the maximum compression principal stress between the two groups aligns well with the two-fold increase in p-MRLC intensity. Together, the prolonged contraction period post amputation and delayed wound closure in *duox* morphants results from reduced tissue rigidity rather than p-MRLC overload at the wound margin (Fig. 5F). Moreover, we demonstrated that the DPI treatment caused over contraction was due to significantly elevated stress at the wound margin through linear elasticity simulation, assuming the DPI-treated group and the DMSO ctrl group share the same Young’s modulus as the *p53* control (Fig. 5G-I). The simulated elevated compression stress at 1000s post-amputation in the DPI-treated group, compared to the DMSO control group, yields a consistent result, with a significantly increased p-MRLC level in the DPI-treated group at 15 mpa (Fig. 3C-E, Fig. 5H-I, and Movie 5). The continued increase in compressive principal stress over time in the DPI-treated group suggests a dramatic rise in force at the wound margin at 1 hpa, which effectively fills the gap in our experimental data (Fig. 5H).

**Figure 5.**
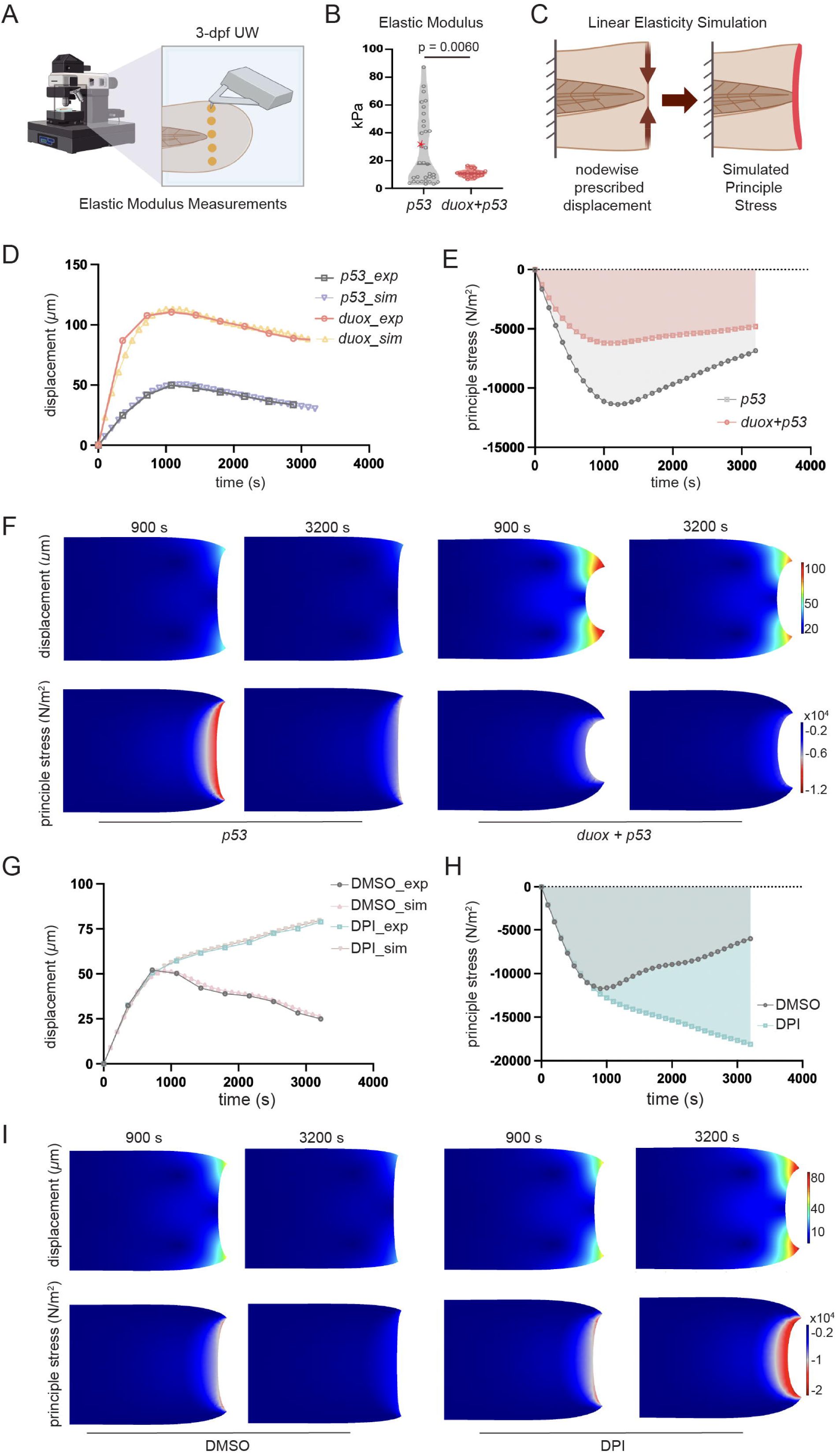
Long-term depletion of ROS reduces tissue stiffness. (A) Schematics of tissue stiffness measurement of MO-injected larvae at 3 dpf. (B) Young’s modulus measurements of the zebrafish tail fin of the *duox* morphant (n = 20) and *p53* ctrl (n = 33) groups. The red asterisk indicates the midrange in the *p53* group. Data are mean ± s.e.m. from at least three independent experiments. (C) Schematics of an *in silico* linear elasticity simulation of the zebrafish tailfin post-amputation. The third principal stress at the wound margin was simulated with a node-wise prescribed displacement derived from experimental data. (D) The displacement of the tailfin edge tip changes over time in both experimental data and *in silco-*fitted data of MO-injected groups. (E) Simulated principal stress at the zebrafish tailfin wound margin over time for the MO-injected groups. (F) Principal stress at the zebrafish tailfin wound margin at 900s and 3200s post-amputation and the corresponding displacement of the MO-injected groups. (G) The displacement of the tailfin edge over time in both experimental data and *in silco-*fitted data of the DPI-treated and DMSO ctrl groups. (H) Simulated principal stress at the zebrafish tailfin wound margin over time for the DPI-treated and DMSO ctrl groups. (I) Principal stress at the zebrafish tailfin wound margin at 900s and 3200s post-amputation, along with the corresponding displacement, of the DPI-treated and DMSO ctrl group. Schematics in (A) and (C) were created using BioRender.com.

## DISCUSSION

Here, we report that ROS counteract over-contraction to relax wounds and promote regeneration. The idea that wound over-contraction impedes wound closure is a significant departure from the prevailing view that contraction is a beneficial process in wound closure. The actomyosin “purse string” drives wound-edge cells to reconnect (Bement, Mandato and Kirsch, no date; Russo *et al*., 2005; Clark *et al*., 2009; Kwon *et al*., 2010; Abreu-Blanco *et al*., 2012). In pinch wound, puncture wound, or laser wound models in *Drosophila*, the actomyosin ring localizes at the wound margin and the rear of wound-adjacent cells (Kwon *et al*., 2010; Abreu-Blanco *et al*., 2012), in response to upstream signaling cues, including altered tension, JNK signaling, and calcium signaling (Kwon *et al*., 2010; Kobb, Zulueta-Coarasa and Fernandez-Gonzalez, 2017). The medial actomyosin network, together with the actomyosin “purse string” anchored by E-cadherin-mediated adherent junctions in the epithelial at the wound edge, collectively contributes to wound contraction and closure in *Drosophila* (Tamada *et al*., 2007; Abreu-Blanco *et al*., 2012; Fernandez-Gonzalez and Zallen, 2013). Furthermore, eliminating branched actin leads to the chiral swirling of linear F-actin filaments, thereby maintaining a stable actomyosin ring and facilitating wound closure (Hui *et al*., 2023). Additionally, in other animal models, such as Xenopus oocyte wounds and human colonic specimens, actomyosin also accumulates at the wound sites to promote wound contraction and closure (Russo *et al*., 2005; Clark *et al*., 2009). Here, we propose an opposing perspective that the contraction process post-amputation is not required for wound closure. This idea is supported by our data, which show that MLCK or ROCK inhibition significantly relaxes the wound without impairing wound closure or subsequent wound regrowth (Fig. 3). A slight reduction in contraction post-amputation may even facilitate wound closure, as the relative distances of nuclei to the wound central plane in the ROCK inhibitor-treated group are slightly smaller than those in the DMSO control group.

The opposing role of wound contraction in zebrafish versus other animal models, particularly in *Drosophila*, may result from multiple reasons. Firstly, it may arise from variations in epithelial structures and different healing strategies. Compared to the monolayer of *Drosophila* epithelium, zebrafish epithelia are composed of two interconnected layers (Gault, Enyedi and Niethammer, 2014). The actomyosin ring is restricted to the suprabasal layer, and cells in the basal layer protrude and crawl to the wound edge. The basal layer may compensate for the loss of the contraction force. Secondly, it may result from distinct wounding geometry. The zebrafish fin fold amputation requires the reconnection of two enveloping epithelial sheets. In contrast, in *Drosophila* laser wounding model, a single epithelium closes the circular wound. In addition, the aquatic environment of the zebrafish may activate strikingly different wound signals compared to drosophila wounding models, such as osmolarity differences, varied mechanical forces at the wound edge, distinct activated growth factors and cytokines, leading to variations of wound healing strategies across species (Kwon *et al*., 2010; Yoo *et al*., 2012; Gault, Enyedi and Niethammer, 2014).

Furthermore, we provide evidence that ROS counteract actomyosin during wound closure. ROS play an essential role in regulating cytoskeleton dynamics through direct and indirect actin modulation (DalleDonne, Milzani and Colombo, 1999; Nimnual, Taylor and Bar-Sagi, 2003; Giannoni *et al*., 2005; Heo and Campbell, 2005; Heo *et al*., 2006; Basuroy *et al*., 2010; Klein *et al*., 2011; Hobbs *et al*., 2014; Steinberg, 2015). DalleDonne reported that ROS directly oxidize actin at Cys^374^, thereby decreasing the speed of actin polymerization and compromising F-actin stability (DalleDonne, Milzani and Colombo, 1999). In addition, ROS indirectly tune actin dynamics and cellular processes by regulating a broad spectrum of proteins, including actin-binding proteins, Rho GTPases, kinases, and protein tyrosine phosphatases (Nimnual, Taylor and Bar-Sagi, 2003; Giannoni *et al*., 2005; Heo and Campbell, 2005; Heo *et al*., 2006; Basuroy *et al*., 2010; Klein *et al*., 2011; Hobbs *et al*., 2014; Steinberg, 2015). ROS oxidize actin-binding proteins, such as myosin II and cofilin, to modulate F-actin stability (Klemke *et al*., 2008; Klein *et al*., 2011; Samstag, John and Wabnitz, 2013). Oxidation of Met394 in myosin II heavy chain decreases ATPase activity and actin contraction levels. Oxidation of cofilin at Cys39, Cys80, Cys139, and Cys147 inhibits actin severing and depolymerization, thereby stabilizing the cytoskeleton. Moreover, redox-sensitive GTPase containing NKCD motif, for instance, Rac1 and RhoA, are susceptible to ROS-induced oxidation. Rac1 oxidization at Cys^18^ accelerates its GDP exchange rate and enhances lamellipodia formation and cell migration (Heo and Campbell, 2005; Hobbs *et al*., 2014). Simultaneously, ROS may accelerate nucleotide dissociation and, therefore, inhibit RhoA activity through the oxidization of Cys^16^ and Cys^20^ in RhoA (Heo *et al*., 2006). Furthermore, ROS orchestra cytoskeleton dynamics by regulating kinases such as focal adhesion kinase (FAK), protein kinase A (PKA), and Src kinases. ROS enhances cell survival rate through FAK phosphorylation at various tyrosine sites, including Y^397^, Y^925^, and Y^577^ (Basuroy *et al*., 2010). ROS-mediated PKA oxidation at Cys^199^ exposes the Thr^197^ in its adjacent activation loop, rendering it more susceptible to phosphatases. The conformational change results in PKA inactivation and reduced cell migration (Steinberg, 2015). Similarly, ROS oxidize Src kinase at Cys245, and Cys^487^ enhances its activation, leading to enhanced cell migration and invasion (Giannoni *et al*., 2005). Additionally, ROS inactivate low molecular weight protein tyrosine phosphatase (LMW-PTP) through the oxidization at Cys^12^. The reduced p190-RhoGAP phosphorylation level downregulates RhoA activity, facilitating cell spreading (Nimnual, Taylor and Bar-Sagi, 2003).

The ability of ROS to stabilize and disassemble F-actin, as observed in previous studies, highlights the complexity of ROS signaling in multicellular tissues (Bahader et al., 2023; Basuroy et al., 2010; Boardman et al., 2004; DalleDonne et al., 1999; Giannoni et al., 2005; Heo et al., 2006; Heo & Campbell, 2005; Hobbs et al., 2014; Nimnual et al., 2003; Steinberg, 2015). In this study, we demonstrate that ROS inhibit the phosphorylation of MRLC to relax the wound, providing a novel context where ROS inhibit actomyosin during wound closure in zebrafish.

We observed smaller epithelial cell areas in *duox* morphants (Fig. 4). This phenotype is likely due to increased cell division, as elevated ROS levels delay the G1-S cell-cycle transition and cell proliferation, resulting in cell enlargement in *Drosophila* eye imaginal disks (Owusu-Ansah *et al*., 2008). We also observed reduced actin abundance in the tailfin and reduced tissue stiffness in *duox* morphants (Fig. 4). This observation is consistent with a previous report that eliminating ROS with NOX inhibitors in zebrafish embryos delays epiboly progression through interference with E-cadherin and cytoskeleton dynamics (Mendieta-Serrano *et al*., 2019; Nikmard *et al*., 2022). Moreover, ROS may mediate the expression of actin and actin-binding protein genes, thereby modulating actin abundance. ROS elimination through NOX1 RNAi silencing significantly reduces SM a-actin expression in rat aortic smooth muscle (RASM) cells, and ROS inhibition downregulates actin cytoskeletal regulatory proteins transcription, including profilin, ARP-3, MRLC, and 14-3-3ζ, in mesangial cells (Clarkson *et al*., 2002; Wang and Sun, 2010).

In summary, reactive oxygen species (ROS) are essential for the cytoskeleton dynamics to drive morphogenesis during development. Their robust production at the wound margin plays a pivotal role in facilitating wound closure. In this work, we demonstrate that ROS eliminate the contraction process following amputation, promoting healing and regrowth. Over-contraction impedes wound closure and subsequent healing and regrowth. These observations are instrumental in understanding and treating mucosal wounds.

## METHODS

### Animals and Stable Line Generation

The zebrafish experiment was conducted in accordance with internationally accepted standards. The Animal Care and Use Protocol was approved by The Purdue Animal Care and Use Committee (PACUC), adhering to the Guidelines for the Use of Zebrafish in the NIH Intramural Research Program (protocol number: 1401001018). MATLAB and the sampsizepwr function were used to calculate the sample sizes required for each experiment based on conservative estimates for the variability in the controls for each type of experiment, with a power of 0.9 (significance level of 0.05) in a two-sample t-test. Data were quantified blindly by an investigator who was not involved in data collection. For generating stable zebrafish lines, microinjections of fish embryos were performed by injecting 1 nl of a mixture containing 25 ng/µl plasmids and 35 ng/µl Tol2 transposase mRNA in an isotonic solution into the cytoplasm of embryos at the one-cell stage. The stable lines were generated as previously described (Deng *et al*., 2011). At least two founders (F0) were obtained for each line. Experiments were performed using F2 larvae produced by F1 fish derived from multiple founders to minimize the artifacts associated with random insertion sites.

### Molecular Cloning

For gene expression in the fish envelope layer of epithelia, a Tol2-krt4 vector that has been reported previously was used in this study (Zhou *et al*., 2018). Target genes were inserted into the Tol2 backbone, which contains the krt4 promoter and SV40 polyA for ubiquitous expression. The MRLC2 gene was cloned from pTK91_GFP-MRLC2 plasmid (Addgene # 46355). The mScarlet fragment used in fish was cloned from our previous plasmid constructs in the lab and inserted into the Tol2-krt4 vector. The In-Fusion cloning (In-Fusion HD Cloning Plus Kit, Clontech) was used for fragment fusion. The incorporated primers were as follows:

mrlc2-F, 5’- TACAAGTGAGCGGCCGCGCGCCACCATGTCGAGCAAACGCGCC -3’;
mrlc2-R, 5’- GGCGACCGGTGGATCCCTTGTACAGCTCGTCCATGCC -3’

### Inhibitor Treatments and ROS Intensity Measurements

Inhibitors were dissolved in dimethyl sulfoxide (DMSO) to prepare stock solutions, which were then further diluted in E3 to achieve working concentrations. DMSO was utilized as vehicle control. DPI and NAC were used as ROS inhibitors, with final concentrations of 100 µM and 200 µM, respectively. A final concentration of 200 µM ROCKIOUT and ML-7 was incorporated to disrupt actin dynamics. Larvae were pretreated with the inhibitor for 30-40 mins before tailfin amputation and maintained in the inhibitor for wound dynamics experiments. Inhibitors were washed out 1 h post-amputation for regeneration assays,

A final concentration of 5 µM pentafluorobenzene sulphonyl fluorescein was utilized for ROS intensity measurements, which was also dissolved in DMSO. Embryos were pretreated with the ROS dye 30 mins before amputation for loading and were washed off with E3 buffer 3 times before amputation. The intensities for each group were measured immediately after amputation and 20 minutes post-amputation with ZEISS Axio Zoom.V16 fluorescent microscope. Emission was excited with a 470/40 band-pass filter and acquired with a 525/50 band-pass filter.

### Morpholino Injections and RT-PCR

MO oligonucleotides (Gene Tools) in Danieau buffer (58 mM NaCl, 0.7 mM KCl, 0.4 mM MgSO_4_, 0.6 mM Ca(NO_3_)_2_, and 5.0 mM Hepes, pH 7.1–7.3) were injected (3 nL) into one-cell-stage embryos. For *duox* knockdown, 100 µM *duox* splice MO1 (5′-AGTGAATTAGAGAAATGCACCTTTT-3′) was used (Niethammer et al., 2009).For *p53* knockdown, 300 µM *p53* MO (5′-GCGCCATTGCTTTGCAAGAATTG-3′) was used (Niethammer *et al*., 2009). For morphotyping of the splicing MOs, RNA was prepared from 3-dpf larvae using the RNeasy kit (Qiagen), and one-step RT-PCR was performed using the SuperScript III One-Step RT-PCR System with Platinum Taq DNA Polymerase kit (Thermo Fisher). The oligonucleotide sequences used for RT-PCR were as follows: duox forward, 5′-CCACAATTACCTGGCCTCCA-3′; duox reverse, 5′-AGGTGTTGCATAAACGCAGG-3 ′.

### Wound Dynamics and Regeneration Assays

The tailfin transaction was carried out on a 3-pdf larva using a razor blade, as described previously (Yoo *et al*., 2012). For wound dynamics assays, the width of the tailfin wound was measured 1 hour post-amputation and normalized to the initial tailfin width immediately after amputation. For regeneration assays, regrowth length was quantified by measuring the distance between the caudal tip of the notochord and the amputated wound at 0, 1, and 2 d after wounding.

### Wound Closure Assay

The embryos were fixed at 0 mpa and 30 mpa in 4% paraformaldehyde (PFA) at 4°C overnight, and the fixed samples were imaged with a Zeiss LSM 880 Inverted Confocal microscope. The Z-stack images were then 3D-rendered in Imaris software to acquire the x-z plane of the amputated wound, and the relative distances between nuclei at the wound edge and the wound central plane were calculated utilizing the codes deposit at GitHub: https://gist.github.com/ding275/7bee9327f85b79c4d9c732969bc6b29b#file-ros-jcs-2025-linear-regression-nucleus-ipynb

### Immunofluorescent Staining

Fish embryos were fixed in 4% PFA overnight at 4°C. After fixation, embryos were permeabilized by proteinase K (10 mg/ml, 15 min for embryos at 3 dpf), and fixed again with 4% PFA for 15 min. Embryos were washed three times with PBS-Tween 20 (0.1%) for 5 minutes each on the rocker, and then blocked in PBS containing 0.3% Triton X-100 and 4% BSA for 3 hours at room temperature. Staining was performed in blocking buffer with p-MRLC primary antibody (1:50 dilution) (Cell Signaling #3675) overnight at 4°C followed by three washes with PBS containing 0.3% Triton X-100 (15 mins each on the rocker) as previously described (Xiong *et al*., 2021). For nuclear staining and F-actin labeling, DAPI and Phalloidin were added to the secondary antibody mixture (Thermo Fisher #A-21236) and stained at room temperature for 4 hours, followed by three washes with PBS with 0.3% Triton X-100 again. Confocal imaging was performed using a Zeiss LSM 880 Inverted Confocal microscope with a Zeiss dry 20x objective (NA 0.8).

### Fluorescent Confocal Microscopy

Fluorescent microscopy imaging of *duox* morphant cell morphology was obtained with a Nikon A1R fluorescent confocal microscope. Larvae at 3 dpf were settled on a glass-bottom dish and mounted with 1% low-melt agarose as described previously. Z-stack imaging was then performed at 28 °C (Barros-Becker *et al*., 2017). The far-red (membrane label) channel and red (myosin light chain) channel were acquired sequentially using a 640-nm laser and a 561-nm laser at 3% power, with a Nikon oil immersion 60x N/A 1.40 objective.

Fluorescent microscopy imaging data from immunofluorescent staining and wound closure assays were acquired using a Zeiss LSM 880 Inverted Confocal microscope with a Zeiss dry 20x N/A 0.8 objective at 2% power. For immunostaining z-stack imaging, the blue, red, and far-red channels were acquired using the 405-nm, 561-nm, and 641-nm lasers, respectively. For wound closure assay z-stack imaging, the green channel was acquired with a 488 nm laser.

### Atomic force microscopy

An Asylum Research MFP3d Bio AFM system (Santa Barbara, CA) was utilized for measuring the effective modulus of *p53* ctrl and *duox* morphants tissue stiffness. A Nanoandmore PPP-CONTAuD-10 probe (Watsonville, CA) with a 450 µm spherical probe and 0.2 N/m nominal stiffness cantilever was used for these experiments. The optical lever sensitivity of the cantilever was calibrated with a static force–displacement curve, which was performed on the part of the polystyrene Petri dish not covered by the embryos. The cantilever stiffness was determined using the thermal tuning method in the E3 medium (Hutter & Bechhoefer, 1993).AFM indentation experiments were performed using morpholino-injected embryos at a constant temperature of 28°C. For imaging, a sufficient amount of E3 buffer was used to maintain embryo viability, and a 200 µl drop of E3 was placed on the AFM cantilever tip to avoid the formation of air bubbles between the cantilever holder and the sample. Force–displacement curves were acquired for 3 biological replicates of MO-injected embryo at 3 positions along the tailfin region. 3 indentations were taken at each position for each fish. The force spectroscopy was conducted using a single loading-unloading cycle, with the trigger threshold set at 50 nN based on cantilever deflection.

To account for stress relaxation, a minimum interval of 45 seconds was maintained between successive indentations within the same embryo. For each measurement location, the force-displacement curve calculated during the cantilever’s approach phase was used to determine the effective modulus. This calculation also incorporated the cantilever’s material characteristics and applied the Hertzian contact model for a spherical indenter interacting with a flat surface (Guo & Akhremitchev, 2006). Based on this model, the force measured during indentation (F) is correlated with cantilever indentation (δ) as described by Johnson (1985).

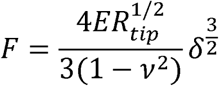

where *v* is the Poisson’s ratio (*v* = .5, assumed for biological samples **need ref), E is the effective elastic modulus, and Rtip is the radius of the spherical cantilever tip. The embryo is assumed to be incompressible with µ = 0.5. The force curves were divided into two regions: one prior to the contact point of the cantilever tip with the tissue and the other after contact. Force data after contact with the embryo was used to calculate the effective modulus.

### Linear Elasticity Simulation

The zebrafish tail fin tissue was modeled as a solid material using the Solid Mechanics Module in COMSOL Multiphysics (Fig. Sup). A 2D finite element model was constructed to represent the tail fin, assuming homogeneous, isotropic material behavior under linear elasticity. The conservation equation undergoing time-varying deformations is derived from Newton’s second law as:

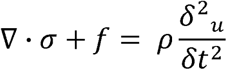

Here, ρ is the (constant) mass density, u is the displacement vector, σ is the Cauchy stress tensor, and f is the body force per unit volume.

The contractile stress was computed based on experimentally measured displacement profiles. For isotropic linear elasticity (Hooke’s law) of a 2D generalization form, the stress σ is related to the strain tensor ε (⛛_*u*_) via:

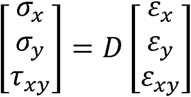

For the isotropic case, D is a function of the Young’s modulus and Poisson’s ratio *v*:

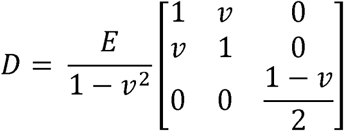

All finite element (FE) simulations were performed in COMSOL using a 2D triangular mesh on a 400□µm□× 500□µm domain to mimic the physiological dimensions observed in experimental imaging. The boundary representing the proximal attachment to the body was constrained to simulate fixed anchorage, ensuring deformation occur only in the distal region. To simulate tissue contraction and acquire mechanical response of the tissue, prescribed node velocity was applied at the distal end of the tail fin, corresponding to the maximum displacement recorded in the experiment for both MO groups.

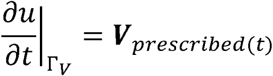

Here, Γ*_v_* represents the relevant boundary in a discrete model. The prescribed node-wise velocity boundary condition was fitted to the Fourier series to reflect the experimentally measured displacements at the wound edge. For different simulation conditions, various functional forms of the boundary velocity are employed, as described below.

DMSO: 0.0145 + 0.0420*cos(t*0.0013) -0.0009*sin(t*0.0013) + 0.0252*cos(2*t*0.0013) + 0.0002*sin(2*t*0.0013) + 0.0081*cos(3*t*0.0013) + 0.0056*sin(3*t*0.0013)

DPI: 0.0227 + 0.0263*cos(t*0.0012) + 0.0087*sin(t*0.0012) + 0.0236*cos(2*t*0.0012) + 0.0066*sin(2*t*0.0012) + 0.0132*cos(3*t*0.0012) -0.0001*sin(3*t*0.0012) + 0.0035*cos(4*t*0.0012) -0.0028*sin(4*t*0.0012)

*p53*: 0.0556*sin(0.0018*t+2.1928) + 0.0064*sin(0.0044*t-0.6985) + 0.0966*sin(0.0008*t+1.5461)

*duox+p53*: 0.0556*sin(0.0018*t+2.1928) + 0.0064*sin(0.0044*t-0.6985) + 0.0966*sin(0.0008*t+1.5461)

To evaluate the compressive stress related to the wound closure, the principal stress was calculated from the local stress state at the node by finding the eigenvalues of Cauchy stress tensor σ:

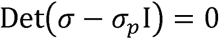

where I is the identity matrix. The corresponding eigenvectors indicate the principal directions, where shear stress components vanish. Negative principal stress (σ*_p_* < 0 corresponds to a compressive normal stress in its principal direction.

### Image Data Analysis

Image analysis was performed with Imaris software and Fiji/ImageJ. For immunofluorescent staining images and optogenetic imaging data, all images at the same time point of an embryo tailfin were projected to 2D for fluorescence intensity measurements. The images were background-subtracted, and the mean intensity values were acquired using Fiji/ImageJ. The myosin distribution and cell morphology data were analyzed both in Fiji/ImageJ and Imaris software. The images were Z-projected, background-subtracted, and median-filtered in Fiji/ImageJ. The perimeters and aspect ratios were measured directly in Imaris software. The wound closure data was 3D-rendered with Imaris (RRID: SCR_007370) software, and nuclei were masked with built-in functions as described.

### Statistical Analysis

Statistical analysis was performed using Prism 6 (GraphPad). An unpaired two-tailed Student’s t-test or ANOVA was used to determine the statistical significance of differences between groups. A P-value of less than 0.05 was considered statistically significant.

### Summary Statement

This study demonstrates that reactive oxygen species regulate wound kinetics, ultimately facilitating efficient wound closure. Acute inhibition of reactive oxygen species with pharmacological inhibitors leads to prolonged wound contraction, likely due to the activation of actomyosin. Meanwhile, long-term reactive oxygen species depletion with morpholino treatment also leads to wound over-contraction, but due to a significant reduction in tissue rigidity and elasticity. Furthermore, this study reveals that contraction is not required for wound healing. Additionally, excessive wound contraction can hinder the wound-healing process. These findings challenge the conventional view that actomyosin ring-mediated contraction provides the primary driving force for wound closure, offering novel insights into the context-dependent role of contraction in wound healing.

## Acknowledgment

We thank the other members of the Emergent Mechanisms in Biology of Robustness Integration and Organization Institute (EMBRIO) for their data analysis support and insightful suggestions.

## Competing interests

The authors declare no competing or financial interests.

## Funding

This work is based upon efforts supported by EMBRIO Institute, contract #2120200, a National Science Foundation (NSF) Biology Integration Institute

## Data availability

All relevant data and resources can be found within the article and its supplementary information.

## Notes

### Competing Interest Statement

The authors have declared no competing interest.

### Summary of Updates

We removed two authors from the manuscript because the data they collected was not included in the submitted version.

## References

Abreu-Blanco, M.T. et al. (2012) ‘Drosophila embryos close epithelial wounds using a combination of cellular protrusions and an actomyosin purse string’, Journal of Cell Science, 125(24), pp. 5984–5997. Available at: 10.1242/jcs.109066.

Barros-Becker, F. et al. (2017) ‘Live imaging reveals distinct modes of neutrophil and macrophage migration within interstitial tissues’, Journal of Cell Science, 130(22), pp. 3801–3808. Available at: 10.1242/jcs.206128.

Basuroy, S. et al. (2010) ‘Hydrogen peroxide activates focal adhesion kinase and c-Src by a phosphatidylinositol 3 kinase-dependent mechanism and promotes cell migration in Caco-2 cell monolayers’. Available at: 10.1152/ajpgi.00368.2009.-Recent.

Bement, W.M., Mandato, C.A. and Kirsch, M.N. (no date) Wound-induced assembly and closure of an actomyosin purse string in Xenopus oocytes. Available at: http://biomednet.com/elecref/0960982200900579.

Clark, A.G. et al. (2009) ‘Integration of Single and Multicellular Wound Responses’, Current Biology, 19(16), pp. 1389–1395. Available at: 10.1016/j.cub.2009.06.044.

Clarkson, M.R. et al. (2002) ‘High Glucose-altered Gene Expression in Mesangial Cells’, Journal of Biological Chemistry, 277(12), pp. 9707–9712. Available at: 10.1074/jbc.M109172200.

DalleDonne, I., Milzani, A. and Colombo, R. (1999) ‘The tert-butyl hydroperoxide-induced oxidation of actin Cys-374 is coupled with structural changes in distant regions of the protein’, Biochemistry, 38(38), pp. 12471–12480. Available at: 10.1021/bi990367k.

David, J.A. et al. (2017) ‘The Nrf2/Keap1/ARE Pathway and Oxidative Stress as a Therapeutic Target in Type II Diabetes Mellitus’, Journal of Diabetes Research, 2017, pp. 1–15. Available at: 10.1155/2017/4826724.

Deng, Q. et al. (2011) ‘Dual Roles for Rac2 in Neutrophil Motility and Active Retention in Zebrafish Hematopoietic Tissue’, Developmental Cell, 21(4), pp. 735–745. Available at: 10.1016/j.devcel.2011.07.013.

Dunnill, C. et al. (2017) ‘Reactive oxygen species (ROS) and wound healing: the functional role of ROS and emerging ROS modulating technologies for augmentation of the healing process’, International wound journal, 14(1), pp. 89–96. Available at: 10.1111/iwj.12557.

Enyedi, B. and Niethammer, P. (2015) ‘Mechanisms of epithelial wound detection’, Trends in cell biology, 25(7), pp. 398–407. Available at: 10.1016/j.tcb.2015.02.007.

Fernandez-Gonzalez, R. and Zallen, J.A. (2013) ‘Wounded cells drive rapid epidermal repair in the early Drosophila embryo’, Molecular Biology of the Cell, 24(20), pp. 3227–3237. Available at: 10.1091/mbc.E13-05-0228.

Finkel, T. (1998) ‘Oxygen radicals and signaling’, Current Opinion in Cell Biology, 10(2), pp. 248–253. Available at: 10.1016/S0955-0674(98)80147-6.

Franco, J.J. et al. (2019) ‘Cellular crowding influences extrusion and proliferation to facilitate epithelial tissue repair’, Molecular Biology of the Cell, 30(16), pp. 1890–1899. Available at: 10.1091/mbc.E18-05-0295.

Fujiwara, T. et al. (2016) ‘Extracellular superoxide dismutase deficiency impairs wound healing in advanced age by reducing neovascularization and fibroblast function’, Experimental Dermatology, 25(3), pp. 206–211. Available at: 10.1111/exd.12909.

Gault, W.J., Enyedi, B. and Niethammer, P. (2014) ‘Osmotic surveillance mediates rapid wound closure through nucleotide release’, The Journal of cell biology, 207(6), pp. 767–782. Available at: 10.1083/jcb.201408049.

Gemberling, M. et al. (2013) ‘The zebrafish as a model for complex tissue regeneration’, Trends in genetics, 29(11), pp. 611–620. Available at: 10.1016/j.tig.2013.07.003.

Giannoni, E. et al. (2005) ‘Intracellular Reactive Oxygen Species Activate Src Tyrosine Kinase during Cell Adhesion and Anchorage-Dependent Cell Growth’, Molecular and Cellular Biology, 25(15), pp. 6391–6403. Available at: 10.1128/MCB.25.15.6391-6403.2005.

Heo, J. et al. (2006) ‘Redox Regulation of RhoA’, Biochemistry, 45(48), pp. 14481–14489. Available at: 10.1021/bi0610101.

Heo, J. and Campbell, S.L. (2005) ‘Mechanism of redox-mediated guanine nucleotide exchange on redox-active Rho GTPases’, Journal of Biological Chemistry, 280(35), pp. 31003–31010. Available at: 10.1074/jbc.M504768200.

Hobbs, G.A. et al. (2014) ‘Rho GTPases, oxidation, and cell redox control’, Small GTPases, 5(2), p. e28579. Available at: 10.4161/sgtp.28579.

Hui, J. et al. (2023) ‘Coordinated efforts of different actin filament populations are needed for optimal cell wound repair’, Molecular Biology of the Cell, 34(3). Available at: 10.1091/mbc.E22-05-0155.

Hutter, J.L. and Bechhoefer, J. (1993) ‘Calibration of atomic-force microscope tips’, Review of Scientific Instruments, 64(7), pp. 1868–1873. Available at: 10.1063/1.1143970.

JIA, N., et al. (2021) ‘Electric Field: A Key Signal in Wound Healing’, Chinese Journal of Plastic and Reconstructive Surgery, 3(2), pp. 95–102. Available at: 10.1016/S2096-6911(21)00090-X.

Kanta, J. (2011) ‘The Role of Hydrogen Peroxide and Other Reactive Oxygen Species in Wound Healing’, *Acta Medica (Hradec Kralove*, Czech Republic), 54(3), pp. 97–101. Available at: 10.14712/18059694.2016.28.

Klein, J.C. et al. (2011) ‘Structural and functional impact of site-directed methionine oxidation in myosin’, Biochemistry, 50(47), pp. 10318–10327. Available at: 10.1021/bi201279u.

Klemke, M. et al. (2008) ‘Oxidation of Cofilin Mediates T Cell Hyporesponsiveness under Oxidative Stress Conditions’, Immunity, 29(3), pp. 404–413. Available at: 10.1016/j.immuni.2008.06.016.

Kobb, A.B., Rothenberg, K.E. and Fernandez-Gonzalez, R. (2019) ‘Actin and myosin dynamics are independent during *Drosophila* embryonic wound repair’, Molecular Biology of the Cell, 30(23), pp. 2901–2912. Available at: 10.1091/mbc.E18-11-0703.

Kobb, A.B., Zulueta-Coarasa, T. and Fernandez-Gonzalez, R. (2017) ‘Tension regulates myosin dynamics during *Drosophila* embryonic wound repair’, Journal of Cell Science, 130(4), pp. 689–696. Available at: 10.1242/jcs.196139.

Kwon, Y.C. et al. (2010) ‘Nonmuscle myosin II localization is regulated by JNK during Drosophila larval wound healing’, Biochemical and Biophysical Research Communications, 393(4), pp. 656–661. Available at: 10.1016/j.bbrc.2010.02.047.

Mateus, R. et al. (2012) ‘In Vivo Cell and Tissue Dynamics Underlying Zebrafish Fin Fold Regeneration’, PLoS ONE, 7(12). Available at: 10.1371/journal.pone.0051766.

Mendieta-Serrano, M.A. et al. (2019) ‘NADPH-Oxidase-derived reactive oxygen species are required for cytoskeletal organization, proper localization of E-cadherin and cell motility during zebrafish epiboly’, Free Radical Biology and Medicine, 130, pp. 82–98. Available at: 10.1016/j.freeradbiomed.2018.10.416.

Mitchison, T.J. et al. (2009) ‘A tissue-scale gradient of hydrogen peroxide mediates rapid wound detection in zebrafish’, Nature (London), 459(7249), pp. 996–999. Available at: 10.1038/nature08119.

Naomi, R. et al. (2021) ‘Zebrafish as a Model System to Study the Mechanism of Cutaneous Wound Healing and Drug Discovery: Advantages and Challenges’, Pharmaceuticals, 14(10), p. 1058. Available at: 10.3390/ph14101058.

Niethammer, P. et al. (2009) ‘A tissue-scale gradient of hydrogen peroxide mediates rapid wound detection in zebrafish’, Nature, 459(7249), pp. 996–999. Available at: 10.1038/nature08119.

Nikmard, F. et al. (2022) ‘The boosting effects of melatonin on the expression of related genes to oocyte maturation and antioxidant pathways: a polycystic ovary syndrome-mouse model’, Journal of Ovarian Research, 15(1), p. 11. Available at: 10.1186/s13048-022-00946-w.

Nimnual, A.S., Taylor, L.J. and Bar-Sagi, D. (2003) ‘Redox-dependent downregulation of Rho by Rac’, Nature Cell Biology, 5(3), pp. 236–241. Available at: 10.1038/ncb938.

de Oliveira, S. et al. (2014) ‘ATP Modulates Acute Inflammation In Vivo through Dual Oxidase 1–Derived H2O2 Production and NF-κB Activation’, The Journal of Immunology, 192(12), pp. 5710–5719. Available at: 10.4049/jimmunol.1302902.

Owusu-Ansah, E. et al. (2008) ‘Distinct mitochondrial retrograde signals control the G1-S cell cycle checkpoint’, Nature Genetics, 40(3), pp. 356–361. Available at: 10.1038/ng.2007.50.

Razzell, W. et al. (2013) ‘Calcium Flashes Orchestrate the Wound Inflammatory Response through DUOX Activation and Hydrogen Peroxide Release’, Current biology, 23(5), pp. 424–429. Available at: 10.1016/j.cub.2013.01.058.

Redd, M.J. et al. (2004) ‘Wound healing and inflammation: embryos reveal the way to perfect repair’, Philosophical Transactions of the Royal Society of London. Series B: Biological Sciences, 359(1445), pp. 777–784. Available at: 10.1098/rstb.2004.1466.

Russo, J.M. et al. (2005) ‘Distinct temporal-spatial roles for rho kinase and myosin light chain kinase in epithelial purse-string wound closure’, Gastroenterology, 128(4), pp. 987–1001. Available at: 10.1053/j.gastro.2005.01.004.

Samstag, Y., John, I. and Wabnitz, G.H. (2013) Cofilin: a redox sensitive mediator of actin dynamics during T-cell activation and migration.

Schäfer, M. and Werner, S. (2008) ‘Oxidative stress in normal and impaired wound repair’, Pharmacological Research, 58(2), pp. 165–171. Available at: 10.1016/J.PHRS.2008.06.004.

Sies, H. et al. (2022) ‘Defining roles of specific reactive oxygen species (ROS) in cell biology and physiology’, Nature reviews. Molecular cell biology, 23(7), pp. 499–515. Available at: 10.1038/s41580-022-00456-z.

Soares, M.A. et al. (2016) ‘Restoration of Nrf2 Signaling Normalizes the Regenerative Niche’, Diabetes, 65(3), pp. 633–646. Available at: 10.2337/db15-0453.

Steiling, H. et al. (1999) ‘Different Types of ROS-Scavenging Enzymes Are Expressed during Cutaneous Wound Repair’, Experimental Cell Research, 247(2), pp. 484–494. Available at: 10.1006/excr.1998.4366.

Steinberg, S.F. (2015) ‘Mechanisms for redox-regulation of protein kinase C’, Frontiers in Pharmacology. Frontiers Research Foundation. Available at: 10.3389/fphar.2015.00128.

Tamada, M. et al. (2007) ‘Two distinct modes of myosin assembly and dynamics during epithelial wound closure’, The Journal of Cell Biology, 176(1), pp. 27–33. Available at: 10.1083/jcb.200609116.

Vatsyayan, R. et al. (2011) ‘Role of 4-hydroxynonenal in epidermal growth factor receptor-mediated signaling in retinal pigment epithelial cells’, Experimental Eye Research, 92(2), pp. 147–154. Available at: 10.1016/j.exer.2010.11.010.

Wang, X. and Sun, Z. (2010) ‘Thyroid hormone induces artery smooth muscle cell proliferation: discovery of a new TRα1 Nox1 pathway’, Journal of Cellular and Molecular Medicine, 14(1– 2), pp. 368–380. Available at: 10.1111/j.1582-4934.2008.00489.x.

Xiong, Z. et al. (2021) ‘In vivo proteomic mapping through GFP-directed proximity-dependent biotin labelling in zebrafish’, eLife, 10. Available at: 10.7554/eLife.64631.

Yoo, S.K. et al. (2012) ‘Early redox, Src family kinase, and calcium signaling integrate wound responses and tissue regeneration in zebrafish’, Journal of Cell Biology, 199(2), pp. 225–234. Available at: 10.1083/jcb.201203154.

Zhou, W. et al. (2018) ‘MicroRNA-223 Suppresses the Canonical NF-κB Pathway in Basal Keratinocytes to Dampen Neutrophilic Inflammation’, Cell Reports, 22(7), pp. 1810–1823. Available at: 10.1016/J.CELREP.2018.01.058.

