## supplemental figure and movie legend for "Reactive Oxygen Species Counteract Wound Contraction and Promote Wound Healing"

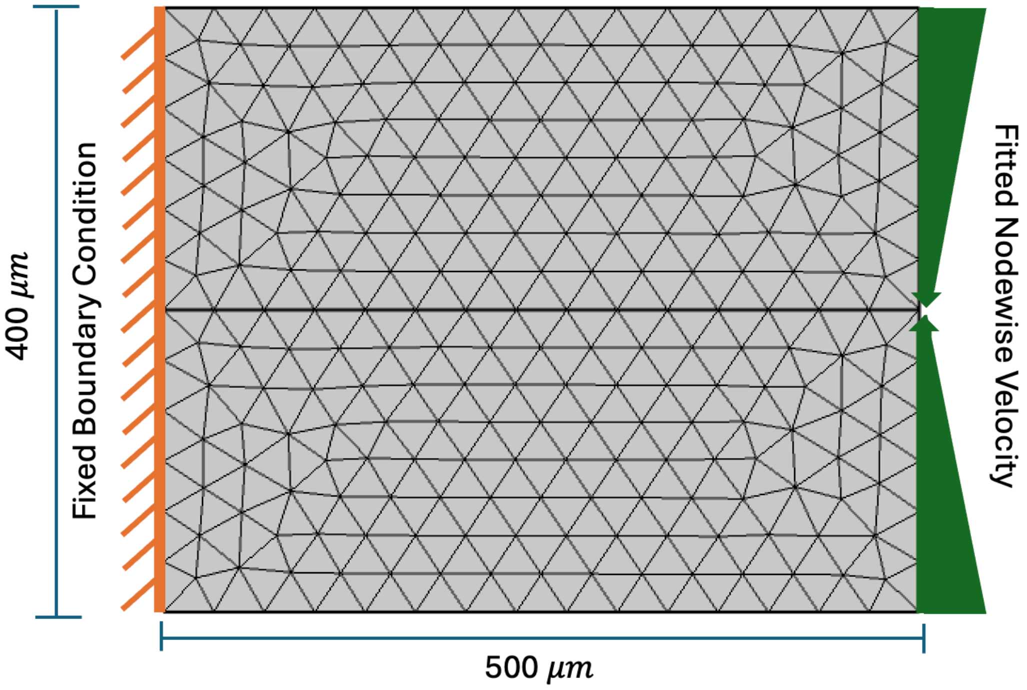


**Figure SI. Finite element (FE) model mesh and Boundary Condition (BC) setting.** Finite element (FE) simulations were performed in COMSOL with a 2D triangular mesh over a 400 μm × 500 μm domain. The material was modeled as linear elastic, with Young's modulus values determined by AFM measurements (*p53*/DMSO/DPI: 45,175 Pa; *duox+p53*: 10,942 Pa) and a Poisson's ratio of 0.49. A fixed constraint was assigned to the left boundary to represent attachment to the main fish body, while the lateral edges were left free to accommodate lateral deformation. At the right boundary, a node-wise velocity boundary condition was prescribed, which was fitted to a Fourier series to reflect the experimentally measured displacements at the cutting edge. Different simulation conditions were implemented by varying the functional form of the boundary velocity.

### Movie legend

**Movie 1. Pharmacological inhibition of ROS reduces wound relaxation post-amputation.**

Time-lapse imaging of zebrafish tailfin wound dynamics of the DPI-treated and DMSO control groups. The imaging started immediately after amputation and continued for an hour, with one-minute intervals. Scale bar: 50 μm.

**Movie 2. ROS inhibition with *duox* knockdown impairs wound relaxation post-amputation.**

Time-lapse imaging of zebrafish tailfin wound dynamics of the *duox* morpholino-injected and p53 control groups. The imaging started immediately after amputation and continued for an hour, with one-minute intervals. Scale bar: 50 μm.

**Movie 3. Pharmacological inhibition of ROCK and MLCK prevents wound contraction.**

Time-lapse imaging of zebrafish tailfin wound dynamics of the ROCKOUT-treated, ML-7-treated, and DMSO control groups. The imaging started immediately after amputation and continued for an hour, with one-minute intervals. Scale bar: 50 μm.

**Movie 4. Linear elasticity simulation of wound dynamics in *duox*-deficient embryos.**

Node-wise prescribed displacement changes over time for MO-injected groups and their simulated corresponding changes in third principal stress over time. The displacement input was fitted to the experimental data from Fig. 1J. The Tissue modulus of the *duox*-injected group and the *p53* ctrl group are 10942 Pa and 45175 Pa, respectively.

**Movie 5. Linear elasticity simulation of wound dynamic upon ROS inhibition.**

Node-wise prescribed displacement changes over time for DPI-treated and DMSO control groups, along with their simulated corresponding third principal stress changes over time. The displacement input was fitted to the experimental data shown in Fig. 1E. The tissue modulus for both groups was 45175 Pa.
